# EGGNet, a generalizable geometric deep learning framework for protein complex pose scoring

**DOI:** 10.1101/2023.03.22.533800

**Authors:** Zichen Wang, Ryan Brand, Jared Adolf-Bryfogle, Jasleen Grewal, Yanjun Qi, Steven A. Combs, Nataliya Golovach, Rebecca Alford, Huzefa Rangwala, Peter M. Clark

**Author notes:** Contributing authors.

## Abstract

Computational prediction of molecule-protein interactions has been key for developing new molecules to interact with a target protein for therapeutics development. Past work includes two independent streams of approaches: (1) predicting protein-protein interactions (PPI) between naturally occurring proteins and (2) predicting the binding affinities between proteins and small molecule ligands (aka drug target interaction, or DTI). Studying the two problems in isolation has limited the ability of these computational models to generalize across the PPI and DTI tasks, both of which ultimately involve non-covalent interactions with a protein target. In this work, we developed an Equivariant Graph of Graphs neural Network (EGGNet), a geometric deep learning framework for molecule-protein binding predictions that can handle three types of molecules for interacting with a target protein: (1) small molecules, (2) synthetic peptides and (3) natural proteins. EGGNet leverages a graph of graphs (GoGs) representation constructed from the molecule structures at atomic-resolution and utilizes a multi-resolution equivariant graph neural network (GNN) to learn from such representations. In addition, EGGNet leverages the underlying biophysics and makes use of both atom- and residue-level interactions, which improve EGGNet’s ability to rank candidate poses from blind docking. EGGNet achieves competitive performance on both a public protein-small molecule binding affinity prediction task (80.2% top-1 success rate on CASF-2016) and an synthetic protein interface prediction task (88.4% AUPR). We envision that the proposed geometric deep learning framework can generalize to many other protein interaction prediction problems, such as binding site prediction and molecular docking, helping accelerate protein engineering and structure-based drug development.

## 1 Introduction

Protein-protein interactions (PPIs) are key to many fundamental biological processes. Proteins mostly perform their functions via non-covalent interactions with three kinds of molecules including proteins, nucleotides and small molecules. The mechanisms of actions for most drugs involve interacting with protein targets to modulate their biological functions and activities. Being able to design drugs, either small molecules or biologics, to selectively bind a protein target with a desirable affinity is critically important for structure-based drug design.

The ability to modulate PPIs with small molecules or synthetic peptides is a core component of therapeutics development. There exist four classes of problems in structure-based drug design for which molecular interactions can be tackled by machine learning (ML) [1], including (i) protein complex property prediction, (ii) binding site/interface identification, (iii) docking (binding pose generation), and (iv) *de novo* design. Property prediction for protein complexes is a task that can also serve as an integral component for other tasks, such as evaluating binding poses generated by docking algorithms or predicting the affinity for computationally designed novel drug candidates. Many ML methods have been developed for two popular groups of property prediction tasks, PPIs and drug-target interactions (DTIs), with the structure of the protein complex provided as input. However, to the best of our knowledge, none of the existing methods can be used across both problems. In this work, we developed a unifying geometric deep learning (GDL) framework for protein complex pose scoring, which encompasses both DTI affinity prediction and PPI interface prediction.

The proposed approach provides a generalizable and scalable representation of macromolecular complexes that can efficiently represent both protein-small molecule complexes and protein-protein complexes. We also desire the representation to be capable of handling synthetic peptides composed of both natural and non-canonical amino acids. 3D graphs are commonly used to represent the 3D structures of individual proteins and protein-small molecule complexes [1]. To increase its generalizbility and scalability, we extend the 3D graph to 3D graph of graphs (GoG) representation.

GoG, also known as Network of Networks (NoN), is a special graph where the nodes in the top-level graph are also graphs [2, 3]. GoGs can be constructed naturally by connecting independent lower-level graphs by pre-defined associations [4], or similarity can be measured by a graph kernel [5]. GoG can also emerge from partitioning a large graph into different subgraphs based on topology or pre-defined rules. Message-passing GNNs have also been extended to operate on both levels of graphs in GoG [4–6]. The GoG data structure has been used to represent drug-drug interaction networks and metabolite networks [4, 6]. However, it has not been applied to model the 3D structures of macromolecules.

In this study, our contributions are two-fold: (i) we developed a featurization procedure to use the GoG data structure to efficiently unify the representations of all types of molecules, including small molecules, intermediate molecules such as peptides, and macromolecules. The 3D GoG representation retains both atomic- and residue-level information of molecular complexes. (ii) We developed EGGNet, an end-to-end geometric deep learning architecture based on an equivariant GNN to learn from the GoG representations of the 3D structures of protein complexes, optionally integrating physics-informed inductive bias to learn atomic-level interactions [7]. Our architecture can be used to predict both DTIs and PPIs. Notably, we achieved the state-of-the-art protein-small molecule binding affinity predictive performance. We further analyzed the effects of different choices of lower-level molecule graph models and evaluated the potential of transfer learning between the DTI and PPI prediction tasks. We also showed that our model improves the outputs of blind docking models.

## 2 Related work

Prior approaches for structure-based prediction of DTIs and PPIs with ML represent 3D structures of protein complexes using three common representations of protein structures based on 3D grids, 3D surfaces, and 3D graphs [1]. Among them, the 3D graph representation is the only one that preserves all of the information from the input protein structure.

For DTI prediction, all three types of representations and some of their combinations have been explored. For instance, KDEEP [8] estimates binding affinities by representing the protein-ligand complex as a 3D grid and learns from this representation using a 3D convolutional neural network (CNN). PotentialNet [9] builds 3D graphs for the protein-ligand complex using their atoms as nodes and chemical bonds and non-covalent interactions as edges, and then leverages message passing graph neural networks (GNN) [10] to learn from the 3D heterogenous graph. PIGNet [7] extended PotentialNet by adding physics-informed inductive bias to a gate-augmented graph attention network (GAT) [11]. The physics-informed inductive bias is encoded by parameterized energy equations calculating a few non-covalent forces based upon the corresponding inter-atomic distances. HoloProt [12] combines the graph and surface representations of proteins. These studies decompose amino acid residues into atoms and bonds, making it difficult to incorporate potentially useful residue-level features for the ML model to learn from, such as embeddings from protein language models [13–16].

3D graph representations that capture the structural information at atomic resolution are computationally more expensive for PPIs than DTIs; therefore, 3D grids are the most prevalent representation of 3D protein complexes in protein-protein interface prediction. For instance, DeepRank [17] is a 3D CNN-based deep learning method that first maps the amino acid residues at the protein-protein interface to a 3D grid centered on the interface. More recently, Deep Local Analysis (DLA) [18] extends such 3D grid representations to an ensemble of 3D grids.

## 3 Methods

### 3.1 Overview of the EGGNet approach

Overall, EGGNet takes the a 3D structure of a protein complex (also known as pose) as the input and makes predictions about the global properties of the pose, such as the binding affinity between two molecules in the protein complex. We refer to this problem as protein complex pose scoring. The input protein complex is comprised of a protein molecule and an interaction partner that takes one of the three types: (1) small molecules, (2) synthetic peptides and (3) natural proteins. EGGNet first converts the input protein complex pose into a GoG to unify the representation of different types of molecules (Section 3.2). Next, EGGNet’s architecture can learn from the GoG representation by coupling equivariant message-passing GNN with different lower-level molecule-to-vector methods (Section 3.3). Further, EGGNet incorporates physics-informed energy inductive biases (Section 3.4).

### 3.2 GoG representation of molecule complexes

Formally, a GoG is a higher-level graph 𝒢 = (𝒱, *ℰ*), where the node set is composed of lower-level graphs 𝒱 = {*G*_1_, *G*_2_, …, *G*_*n*_ }. The lower-level graph is used to represent the molecule’s building blocks at atomic resolution, hence, *G*_*i*_ = (*V*_*i*_, *E*_*i*_) is a graph with atoms as nodes *atom* ∈*V* and covalent bonds as edges *bond* ∈ *E*. For convenience we represent all notations used in this paper in Table 1.

**Table 1.**
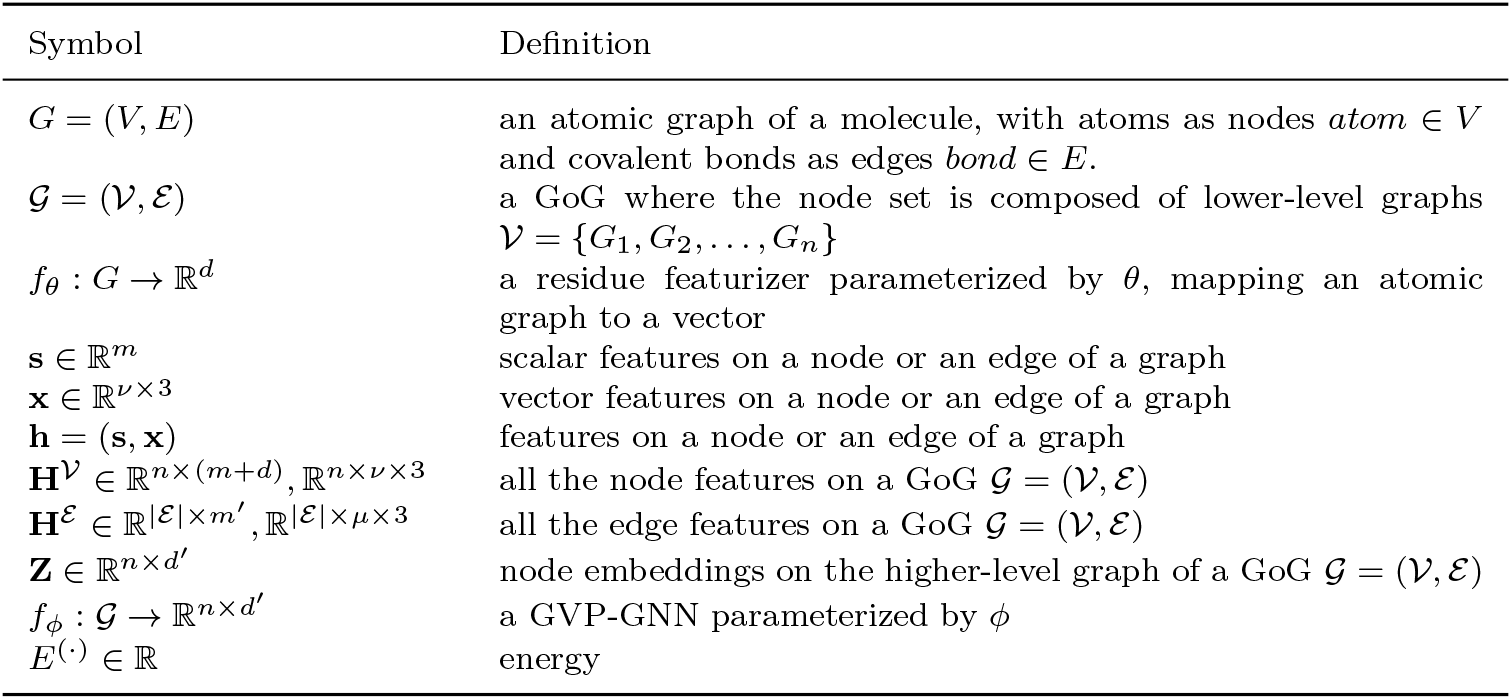
Table of notations.

Given a molecular structure, one can directly construct the atomic resolution graph for the entire molecule by connecting atoms with covalent bonds. We denote this graph as *G* = (*V, E*). To construct a GoG from *G*, we first perform edge-cut graph partitioning to partition *V* into disjoint subsets *V*_1_∪ … ∪*V*_*n*_ = *V* of all the atoms *v*_*u*_ ∈ *V* in graph *G*, resulting in *n* subgraphs, each of which corresponds to a lower-level graph *G*_*i*_ = (*V*_*i*_, *E*_*i*_). We perform the graph partition by cutting the molecular graph at peptide bonds (Fig 1). Next, we construct the higher-level graph 𝒢 = (𝒱, ℰ) using k-nearest neighbor (kNN) algorithm with *k* = 30. We measure the Euclidean distance between the lower-level graphs using the C-alpha or the geometric centroids of heavy atoms, if the subgraph is not an amino acid residue.

**Fig. 1.**
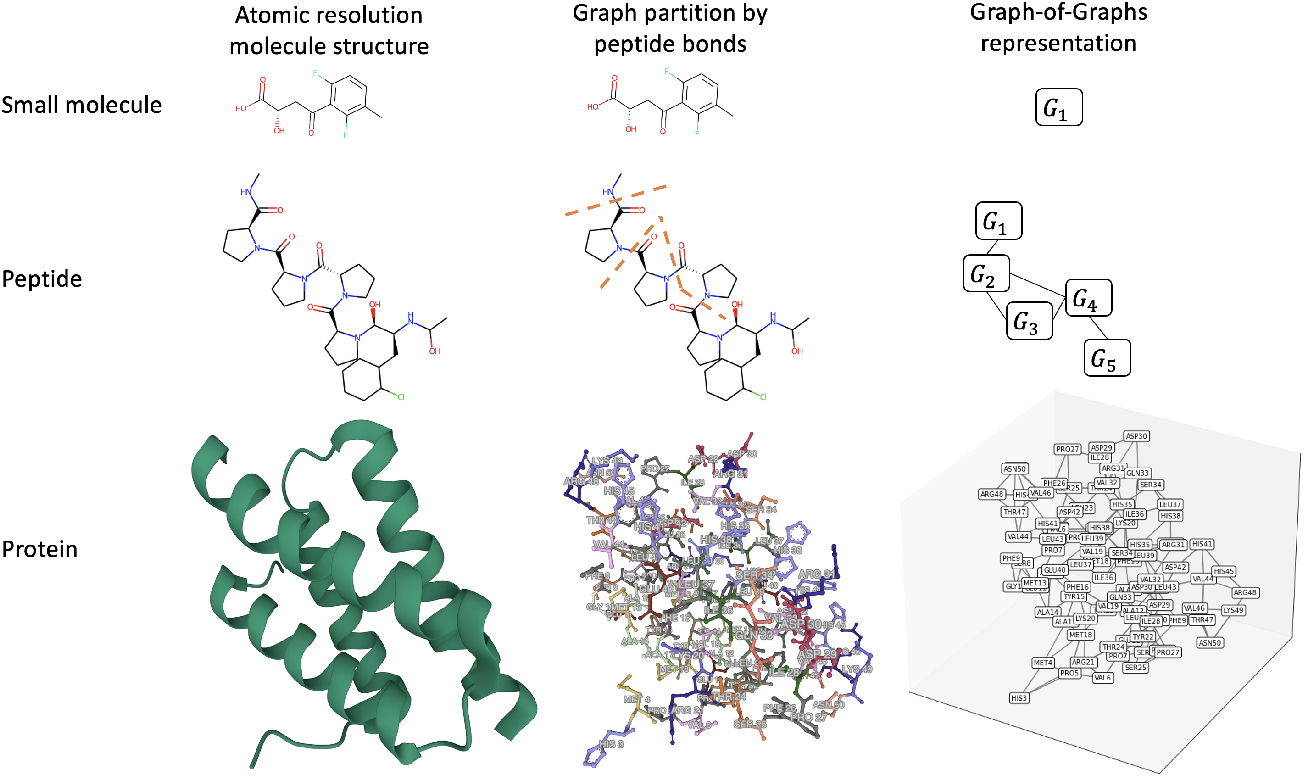
Graph-of-Graphs (GoGs) representation of molecular structures. To construct GoG from molecular structure (left column), we first perform graph partition by cutting peptide bonds (middle column), then connect subgraphs using kNN, leading to a higher-level graphs (right column). The figure shows how GoG representation can generalize to small molecules, peptides with non-canonical amino acid residues, and proteins

With this procedure, we can unify the representations of small molecules and macromolecules: small molecules is a special case of GoG where the higher-level graph is composed of a single node (i.e. 𝒢 = (𝒱= {*G*_1_}, ℰ)). The GoG representation has the advantage over one-hot encoding with fixed vocabulary commonly used in protein language models [13–16], because it can generalize to non-canonical amino acids such as penicillamine used in synthetic peptides. We also use the same GoG construct to represent a protein in complex with other molecules, including small molecules, peptides, or proteins. To do that, we simply apply the edge-cutting graph partition procedure to every polymer chain to derive the lower-level graphs and then construct the higher-level graph by using kNN to connect spatially proximal lower-level graphs (Fig 2).

**Fig. 2.**
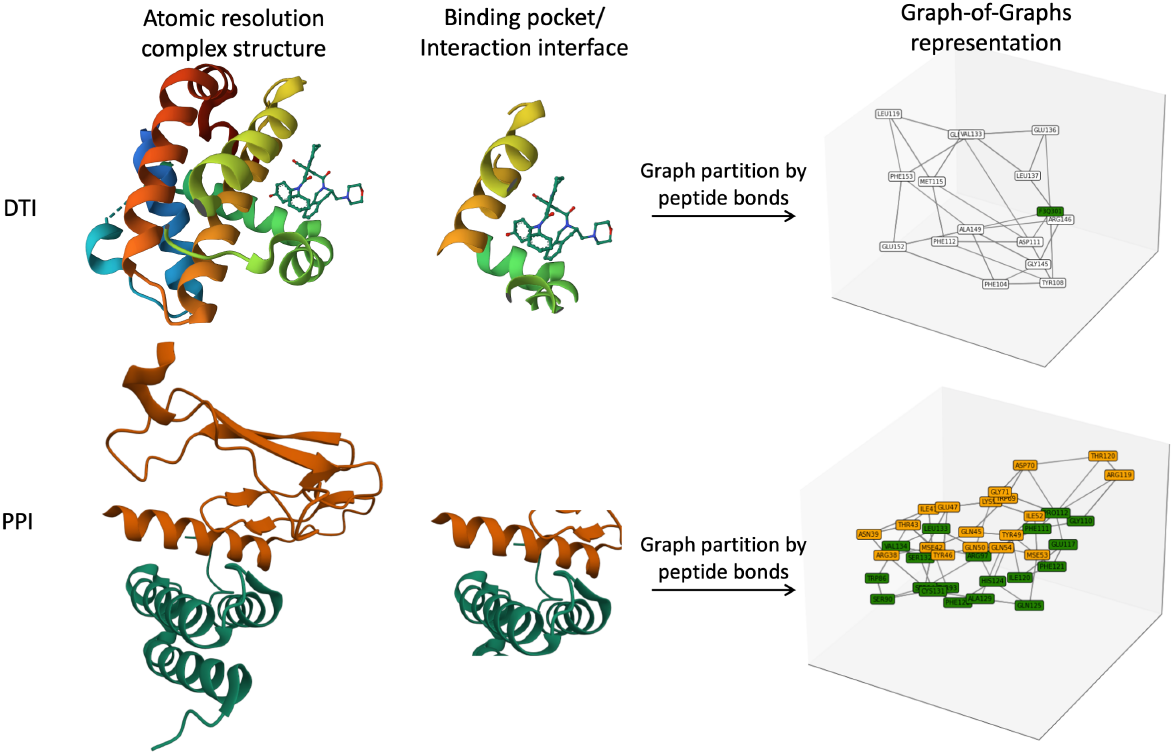
Graph-of-Graphs (GoGs) representation of protein complexes. To construct GoG from the structures of protein complexes (left column), we first identify the binding pocket or interaction interface between molecules (middle column), then perform graph partition by cutting peptide bonds followed by connecting subgraphs using kNN, leading to a higher-level graphs (right column). The nodes in the higher-level graphs of GoGs are colored by the origin of the molecules. The figure shows how GoG representation can generalize to drug-target interaction (DTI) and protein-protein interaction (PPI).

### 3.3 Model architectures for protein complex pose scoring

This section describes the novel deep learning architectures for protein complex pose scoring. The overall architectures for EGGNet are shown in Fig.3

**Fig. 3.**
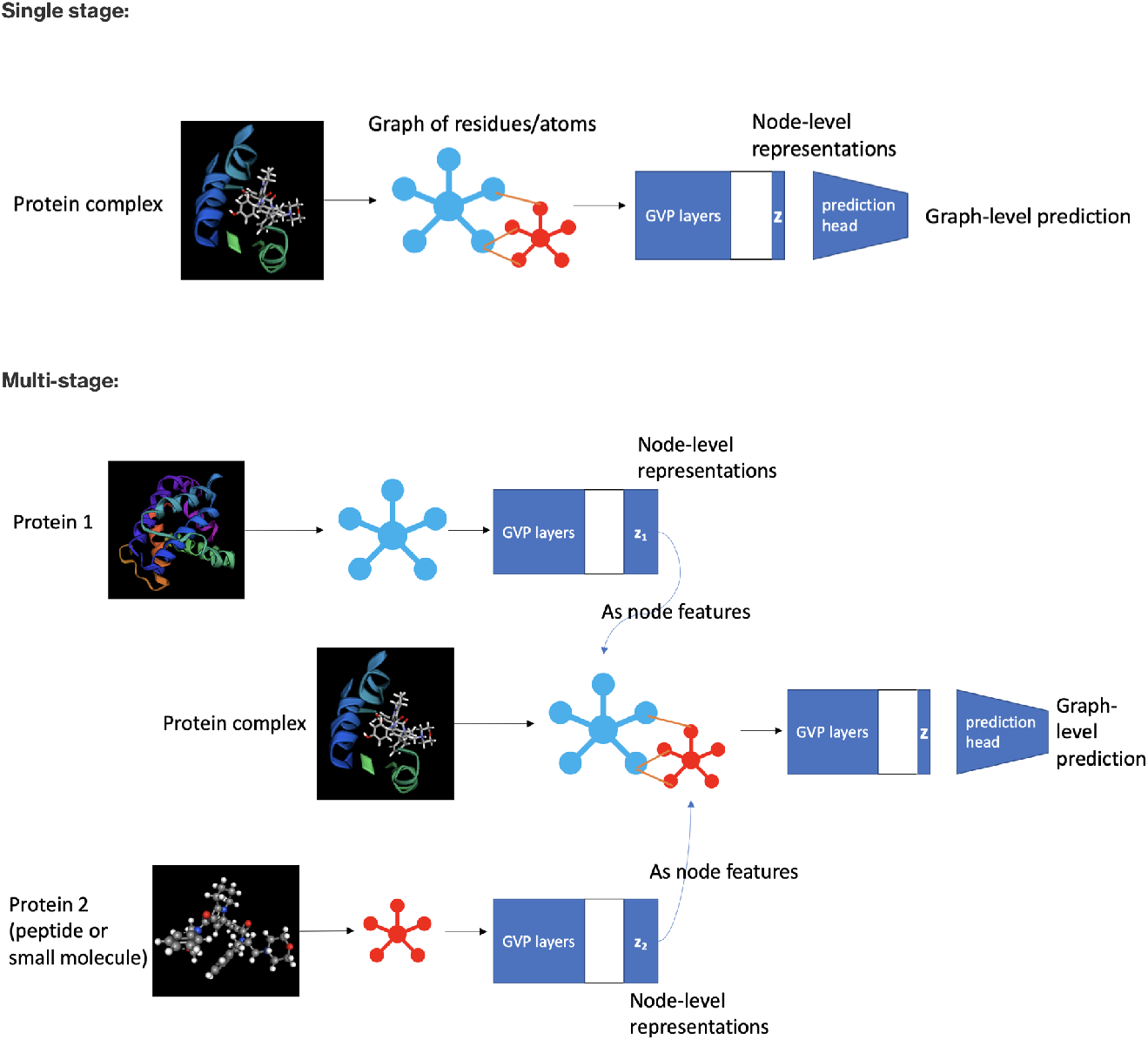
Model architectures. Top: single-stage EGGNet. Bottom: multi-stage EGGNet.

We formulate the protein complex pose scoring problem as a supervised graph-level prediction task. The goal of this task is to learn a function *ŷ* = *f* (𝒢) mapping a GoG 𝒢 representing the protein complex’s structure (pose) to a scalar value *y*, representing the global property of the pose such as the binding affinity between the molecules.

Let 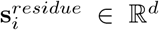 be the feature vector extracted from the lower-level graph/residue. There are many possible choices of the residue featurizer *f*_*θ*_ : *G* →ℝ^*d*^ including 1) chemical fingerprints such as MACCS and ECFP/Morgan [19], 2) GNN models trained on small molecule graphs, and 3) language models trained on the SMILES strings of small molecules such as MolT5 [20].

We denote the node features on higher-level graph as 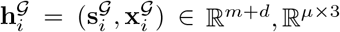, for simplicity, we denote the node and edge features as **H**^**𝒱**^ ∈ℝ^*n×*(*m*+*d*)^, ℝ^*n×ν×*3^ and **Hℰ** ∈ ℝ^|*ℰ*|*×m′*^, ℝ^| *ℰ*|*×μ×*3^, respectively.

GVP-GNN [21] is a *SE*(3)-equivariant GNN in which all node and edge embeddings are tuples (**s, x**) of scalar features **s** ∈ℝ^*m*^ and geometric vector features **x** ∈ ℝ^*ν×*3^. GVP-GNN operates on the node and edge features from the higher-level graph:

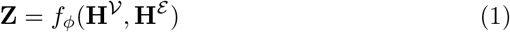

Next, we use a readout network/prediction head to take the learned equivariant node representations **Z** ∈ ℝ^*n×d′*^ to make graph-level prediction:

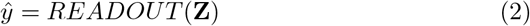

We also design a multi-stage variant of our model architecture to have two GVP-GNNs to learn from two GoGs representing the two interacting molecules 𝒢_1_ = (𝒱_1_, *ℰ*_1_), which are two subgraphs of the complex GoG 𝒢 = (𝒱, *ℰ*), where 𝒱 = 𝒱_1_ ∪ 𝒱_2_ and ℰ = ℰ_1_ ∪ ℰ_2_ ∪ ℰ_*int*_. ℰ_*int*_ denotes the edges representing intermolecular interactions.

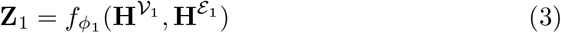

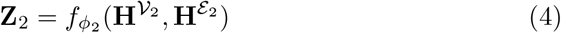

The second stage GVP-GNN then computes node representations:

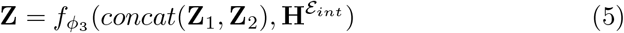

### 3.4 Objective functions and model training

We use mean squared error (MSE) and binary cross-entropy for binding affinity regression and binary interaction prediction tasks, respectively. Additionally, we adopted the physics-informed energy inductive biases from PIGNet [7]. Briefly, we use an energy decoder as the readout function to approximate four types of non-covalent interaction energies (van der Waals interactions, hydrogen bonds, metal-ligand interactions, and hydrophobic interactions) from a protein complex using parameterized equations.

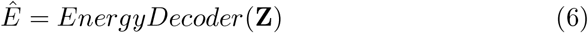

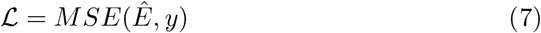

All the parameterized equations takes the atom-level representation **z**_*i*_, **z**_*j*_ ∈ ℝ^*d′*^ as inputs. For instance, van der Waals interaction takes the following form:

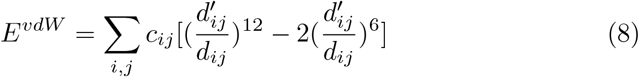

where *d*_*ij*_ = ∥ **z**_*i*_ **z**_*j*_ ∥ and 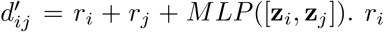 denotes the radii or the *i*-th node.

We also decompose the energy from the whole protein complex to intramolecular (monomer) and intermolecular energies *E* = *E*^(1)^ + *E*^(2)^ + *E*^*int*^ as novel physics inductive bias.

We use stochastic gradient descent (SGD) to learn the parameters *ϕ*, and optionally the residue featurizer parameter *θ*, to optimize the objective function. When using SGD to learn both *ϕ* and *θ*, we essentially allow the end-to-end training of the lower-level residue featurizer and the higher-level GVP-GNN. Similar end-to-end joint training of models operating at different data modalities has also been used for protein function prediction [22].

### 3.5 Datasets and experiments

We use PDBbind [23], ProtCID [24], MANY [25], and DC [26] datasets for training and validation of our models. These datasets cover two otherwise distinctive tasks, DTI and PPI prediction that can be unified by our approach. For the DTI binding affinity regression task, we used the same setup of the PDBbind and CASF-2016 [27] datasets as the PIGNet study [7]. Briefly, the PDBbind 2019 refined set provides quantified binding affinity data (in pKd, where Kd is the experimentally measured dissociation constant) and corresponding structure of protein–ligand complexes deposited in the protein data bank (PDB) [28]. We used 4514 samples for the training set, which is the PDBbind 2019 refined set after removing the redundant samples from the cores set of PDBbind 2016. We randomly sample 20% from the training set as the validation set for early-stopping. For hold-out evaluation, we used the CASF-2016 benchmark dataset, which originated from the PDBbind 2016 core set. We used 283 samples for the scoring and ranking, 22,340 samples for the docking benchmark. All protein-ligand complexes were processed, to focus on the pocket-ligand structure, which removes amino acid residues whose minimum distance between the ligand is greater than 5Å to remove amino acid residues distant to the protein’s binding pocket. We adopt the evaluation metrics recommended in the CASF-2016 dataset [27]. Briefly, we calculate the Spearman’s correlation coefficient *rho* as the evaluate metric for the ranking setting in CASF-2016. For the docking setting, we used the predicted binding affinities to rank candidate poses from each protein-ligand pair, then define a success docking if the top-1 ranked pose has the lowest root-mean-square deviation (RMSD) among candidate poses (decoys). We then compute the top-1 success rate across all protein-ligand pairs as the fraction of successful docking from all docking experiments in the CASF-2106 benchmark.

ProtCID dataset [24] is used for the binary binding classification task. Here, the aim is to distinguish physiological binders with non-physiological ones. Physiological binders were defined as those homo- and hetero-dimers which had at least 5 crystal forms. If an interface is only seen in one crystal form of this UniProt ID in addition to not being in any common cluster, then it is likely to be non-physiological. For data preprocessing, we remove amino acid residues whose C-beta distance is greater than 6Å to any amino acids from the protein chain of the binding partner. We also removed large interfaces with more than 10^5^ atoms, leading to 15,736 and 3,889 protein interfaces for training set and hold-out test set, respectively. The train/test split was performed by sequence clusters resulting from MMseq2 [29] with identity cutoff of 30%.

Similarly, the MANY [25] and the DC [26] datasets contain physiological and non-physiological (crystal) dimers in balanced proportions. We download the datasets from the SBGrid data repository https://data.sbgrid.org/dataset/843/ and follow the same experimental setup as in [17, 30]. Briefly, we train our model using 80% of the MANY dataset, with 20% as the validation set and test the model performance on the hold-out DC dataset. For data preprocessing, we experimented a range of cropping threshold from 6 to 14Å to when removing amino acids residues outside of the interfaces (Fig A1).

We evaluate the model performance on the binary classification task using the area under the receiver operating characteristic curve (AUROC) and the area under the precision-recall curve (AUPR).

For EGGNet models trained on the above datasets, we set the GVP-GNN to have 3 GVP convolutional layers with node and edge hidden dimensions to be (200, 32) and (64, 2), respectively. We used the following hyperparameters for model training: learning rate of 1e-4, per-GPU batch size of 16. To avoid overfitting, we employed an early stopping criterion with patience=50 epochs and trained for a maximum of 1000 epochs. ADAM [31] optimizer with *β*_1_ = 0.9, *β*_2_ = 0.999 is used for optimizing the learnable parameters. All models are implemented using the Pytorch deep learning library [32] and training are performed using Pytorch-Lightning library with 16-bit mixed precision training using 4 NVIDIA V100 GPUs with 16GB of memory each on Amazon SageMaker.

## 4 Experimental Results

### 4.1 GoG representation of molecules allows flexible transfer learning of molecule-to-vector (Mol2Vec) methods

In this work, we leverage the GoG data structure to represent the 3D structures of small- and macro- molecules. Macro-molecules, such as proteins and nucleic acids, are composed of small molecule building blocks (amino acids and nucleotides) connected by covalent bonds. Inspired by this observation, we view a macro-molecule as a graph of small molecule residues, which can be represented by graph of atoms connected by chemical bonds. The GoG representation of molecules provides the flexibility in modeling macro-molecules with out-of-vocabulary residues such as post-translational modifications (PTMs) and non-canonical amino acids. It also unifies the chemical space of small- and macro-molecules by applying Mol2Vec methods to the lower-level graphs.

We first demonstrate that EGGNet is able to take advantage of any Mol2Vec representation approaches on the GoG representation of protein in complex with small molecule ligands. We experimented with three classes of Mol2Vec approaches: (i) chemical fingerprint approaches including MACCS keys and Morgan fingerprints [19], (ii) GNNs pretrained on a small molecule chemical library using structures and properties, and (iii) language models pretrained on SMILES strings of small molecules. We evaluated the performance of single-stage EGGNet with different Mol2Vec methods on the PDBbind/CASF-2016 binding affinity regression task. Our results show that Mol2Vec methods based on deep learning such as Graph Isomorphism Network (GIN) [33] and MolT5 [20] achieved better performance than chemical fingerprints (Table 2) by 0.9%. We also observe that EGGNet with the MolT5 model achieves an improved docking success rate than with GIN by 1.3%, suggesting the more powerful pretrained Mol2Vec model with can further boost the overall model performance. However, MolT5 models (MolT5-small: ∼77M parameters, MolT5-base: ∼250M parameters) are significantly larger than GIN (∼1.8M parameters). As such, we use GIN for later experimentation due to training efficiency.

**Table 2.**
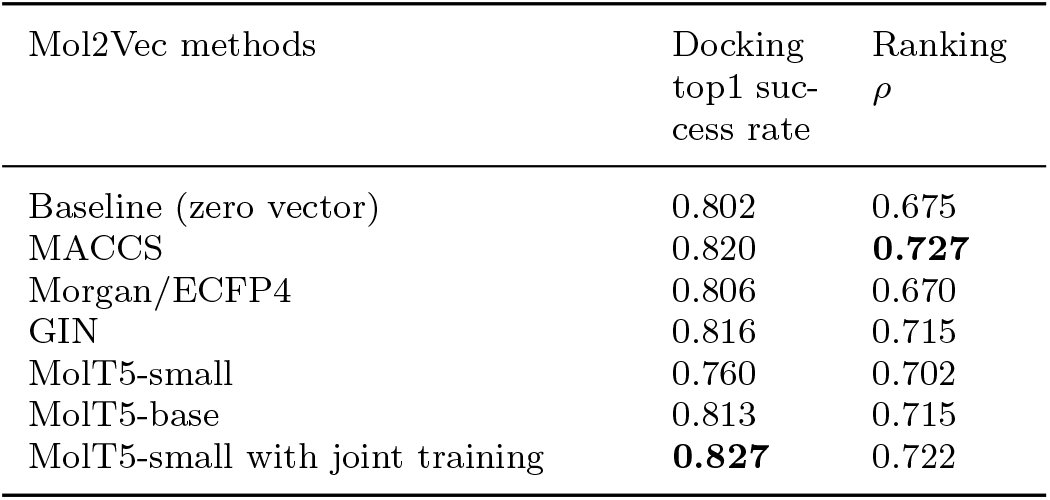
Residue-level representation models improve binding affinity prediction. The following experiments use GVP backbone with energy inductive biases and identical training hyperparameters and setups, except for the residue-level model.

### 4.2 Performance of EGGNet on protein-molecule binding prediction benchmarks

Next, we evaluated EGGNet on two distinctive protein complex pose scoring tasks: (i) the protein-small molecule binding affinity prediction and (ii) proteinprotein interface classification.

#### 4.2.1 Protein small molecule binding affinity regression

On the binding affinity prediction task, we train our model on PDBbind and evaluated on CASF-2016 ranking and docking benchmarks. Our results show that EGGNet with joint training and energy inductive biases outperforms PIGNet [7] in the same setting on both docking and ranking benchmarks (Table 3). We found our best performing model on the docking setting achieved 80.2% top 1 success rate over 77.4% reported in PIGNet.

**Table 3.** Model performances on the CASF-2016 binding affinity regression task.

| Model | Lower-level GNN | Model back-bone | Energy inductive bias | Docking top1 success rate | Ranking $\rho$ |
| --- | --- | --- | --- | --- | --- |
| Ours | GIN | GVP | None | 0.230±0.005 | 0.653±0.036 |
| | GIN | MS-GVP | None | 0.148±0.019 | 0.566±0.031 |
| | GIN joint training | GVP | None | 0.205±0.009 | 0.715±0.019 |
| | GIN joint training | MS-GVP | None | 0.200±0.023 | 0.690±0.009 |
| | GIN | GVP | $E$ | 0.759±0.020 | 0.719±0.025 |
| | GIN | MS-GVP | $E$ | **0.802±0.020** | 0.658±0.032 |
| | GIN joint training | GVP | $E$ | 0.779±0.021 | **0.765±0.023** |
| | GIN joint training | MS-GVP | $E$ | 0.788±0.007 | 0.760±0.015 |
| 3D GNN | None | GNN | $E$ | 0.299 | 0.604 |
| PIGNet | None | GNN | $E$ | 0.774 | 0.672 |

Interestingly, we found the joint training of the GNNs at both the lower-level and higher-level graphs of the GoG improves the models’ performance on the ranking task, while slightly decreasing performance on the docking task. The ranking task examines a model’s ability to generalize to different combinations of different protein-ligand pairs, which corresponds to different composition and geometry of the GoGs. Meanwhile, the docking task probes the models’ ability to rank candidate poses of the same protein-ligand pair, which corresponds to GoGs with the same composition of lower-level graphs but different geometries. Therefore, we reason that the model with the ability to adjust the embeddings of the lower-level residue representations are only beneficial to tasks that prioritize in distinguishing the composition, rather than the geometry of the GoGs. This also explains the observation that the multi-stage variant of EGGNet is more performant on the docking setting (Table 3). The multi-stage variant has more learnable parameters that model the inter-molecular geometries.

We also confirmed the findings from Moon *et al* [7] that using the non-covalent interaction energy as inductive bias for the deep learning model can significantly improve binding affinity prediction. Without the energy inductive bias, the model can only achieve 22.9% top-1 success rate, compared to 75.8% with the energy inductive bias (Table 3). As the energy *E* was calculated for the entire protein-ligand complex, we hypothesized that the inductive bias is more informative to the model if we decompose it to *E* = *E*^(1)^ + *E*^(2)^ + *E*^*int*^. That is, for 2-molecule complex, the non-covalent interaction energy can be decomposed as the sum of intra-molecule energy and inter-molecule energy. However, our results on the PDBbind/CASF-2016 experiments do not support this hypothesis (Table A1). This is probably due to the inaccurate estimation of the protein’s energy *E*^(1)^ because we used only the binding pocket rather than the entire protein structure to construct the GoG to make the training memory efficient.

#### 4.2.2 Protein-protein interface classification

Next, we evaluated EGGNet on another common type of protein complex pose prediction task to distinguish whether a protein-protein interaction interface is physiological or an artifact of the crystallization process. This task can be useful to deduce solution-state quaternary states (dimers, trimers, etc.) and/or to identify novel functions of protein families that were originally deemed crystal artifacts [34]. In addition, models trained for this task could be fine-tuned for detection of binders and non/weak – binders for protein interface design.

We trained and validated our models on the ProtCID [24] dataset. We found different variants of our models are able to significantly outperform CAMP [35], which only takes the sequence information of the protein pairs (Table 4). We also found that the joint training of GNNs at different levels of the GoGs improve the predictive performance over freezing the lower-level GIN by at least 5.8% in AUROC.

**Table 4.** Binary protein-protein interaction prediction on ProtCID.

| Model | Lower-level GNN | Model back-bone | AUROC | AUPR |
| --- | --- | --- | --- | --- |
| Ours | GIN | GVP | 0.673 | 0.858 |
|  | GIN | MS-GVP | 0.671 | 0.857 |
|  | GIN joint training | GVP | 0.734 | 0.875 |
|  | GIN joint training | MS-GVP | **0.745** | **0.890** |
| CAMP | None | CNN | 0.522 | 0.800 |

Next, we evaluated whether the GoG representation leveraged by EGGNet is comparable to other types of representation of protein-protein interfaces. We trained and evaluated EGGNet on the publicly available MANY/DC datasets for protein-protein interface classification (Table 5). We found our model outperforms traditional ML approaches including PISA [36] and PRODIGY-crystal [37, 38] by 8.8% and 1.9%, respectively. Both PISA and PRODIGY-crystal operates on tabular features computed from the interfaces. EGGNet is also competitive with DeepRank-GNN [30], a recently developed GNN-based method. We noted approaches using graph representations underperform DeepRank [17], which represents the interfaces as 3D grids. It is worth pointing out that the featurization process and representation of EGGNet is generalizable to any molecular interaction interfaces, whereas competing methods including DeepRank [17] and DeepRank-GNN [30] rely on features specific to proteins such as pre-computed position-specific scoring matrices (PSSM).

**Table 5.** Protein interface classification performances of models trained on the MANY dataset and tested on the DC dataset. Underlined accuracy scores are reported in [30]

| Model | Interface representation | AUROC | AUPR | Accuracy |
| --- | --- | --- | --- | --- |
| PRODIGY-crystal | tabular | - | - | <u>0.74</u> |
| PISA | tabular | - | - | <u>0.79</u> |
| DeepRank | 3D grid | - | - | <u>**0.86**</u> |
| DeepRank-GNN<sup>1</sup> | 3D graph | 0.865 | **0.871** | 0.820 |
| EGGNet+GIN joint training | 3D GoG | 0.861 | 0.826 | 0.805 |
| EGGNet+MolT5-small joint training | 3D GoG | **0.869** | 0.863 | 0.805 |

<sup>1</sup>The metrics are computed using the model predictions provided by [30]:
https://github.com/DeepRank/Deeprank-GNN

The evaluation of interface classification is similar to the ranking setting of the PDBbind/CASF-2016, where the model is tasked to classify different protein-protein interaction pairs rather than the same protein-protein pair with different poses. Therefore, we observed that the energy inductive bias is not useful in improving the binary classification performance (Table A2).

Thanks to the uniform GoG representation of the protein complexes, our model architectures used for the protein-protein interface prediction and the protein-small molecule binding affinity prediction are identical. We next evaluate if a model trained on the binding affinity prediction task can improve protein-protein interface prediction via transfer learning. However, our results show that transfer learning from the model trained to predict small molecule binding affinity does not help with this task compared to a model trained from scratch (Table A3). We speculate that this is due to the distributions of the input GoGs and the labels between these datasets are disparate.

### 4.3 EGGNet improves ranking of docking poses from DiffDock

Although our model is developed to predict the property of a protein complex using the 3D structure as input, we assess whether our model is helpful to tasks beyond property prediction, such as blind docking, another important task in structural-based drug discovery. Binding affinity prediction is the next step for blind docking, which generates poses for how a drug candidate would bind to a protein target.

We applied our model to predict the binding affinity on candidate poses generated by the state-of-the-art blind docking algorithm DiffDock [39]. Diff-Dock is a composed of a generative model to generate candidate poses for a small molecule with respect to a protein target, and a confidence model to rank the quality of each pose. We compared the rankings of candidate poses by DiffDock’s confidence model, the predicted binding affinity from our model, as well as the combination of both using rank product (Table 6).

**Table 6.**
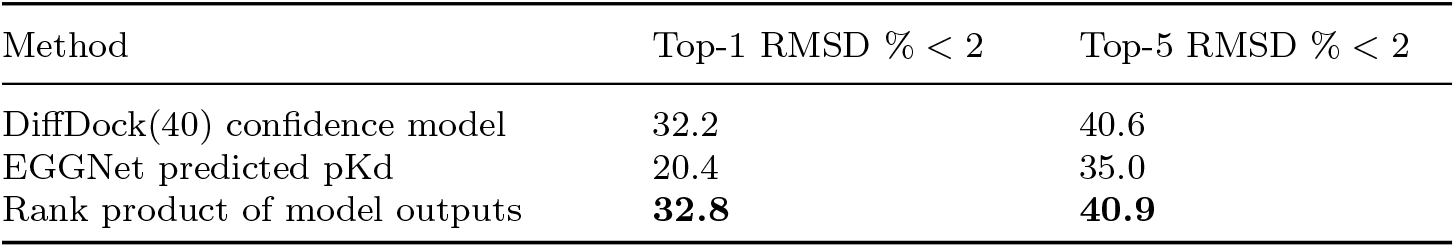
Binding affinity predicted by our model improves ranking of DiffDock pose candidates. Blind docking experiments are performed on the test set of PDBBind v2020.

We found that although the predicted binding affinity alone from our model is not as good as prioritizing the best poses, combining the predicted binding affinity with DiffDock’s confidence measures slightly improves the prioritization of poses, suggesting predicted binding affinity are helpful information in identifying the experimentally validated poses of protein-small molecule complexes. We also envision using our model architecture to jointly learn the RMSD and binding affinity of each candidate pose to further improve generative ML models of blind docking.

## 5 Conclusion

In this work, we first developed a biologically inspired data structure GoG to represent the structures of molecules and molecular complexes. We construct GoGs by applying edge cutting graph partition on the atomic graphs of molecules, which is inspired by the fact that macromolecules are composed of small molecule residues connected by common covalent bonds such as peptide bonds for proteins and phosphodiester bonds for nucleic acids. Next, we developed EGGNet, an equivariant graph neural network to learn from the GoG representations of molecular complexes to predict their properties. The unifying representation allows EGGNet to perform both DTI and PPI prediction tasks, achieving competitive performances. We also demonstrated the value of EGGNet in improving results from blind docking.

## Acknowledgments

We thank Sandeep Somani from Janssen and Tong Wang, Nkechinyere Agu from AWS for insightful discussions.

## Data availability

The PDBBind/CASF-2016 data can be downloaded from https://zenodo.org/record/6047984. The MANY/DC datasets can be downloaded from the SBGrid data repository https://data.sbgrid.org/dataset/843/.

## Code availability

The source code for this study are available for research and non-commercial use at: https://github.com/aws-samples/eggnet-equivariant-graph-of-graph-neural-network

## Appendix A Appendix

**Table A1.** Decomposing energy terms doesn’t improve the model performances on the CASF-2016 binding affinity regression task. *E* = [*E*^*vdW*^, *E*^*hbond*^, *E*^*metal*^, *E*^*hydrophobic*^]

| Energy inductive bias | Docking top1 success rate | Ranking $\rho$ |
| --- | --- | --- |
| $E$ | **0.827** | **0.722** |
| $\mathbf{W}([E^{(1)}, E^{(2)}, E^{int}])$ | 0.583 | 0.638 |
| $\mathbf{W}(\text{BatchNorm}([E^{(1)}, E^{(2)}, E^{int}]))$ | 0.442 | 0.677 |
| $\mathbf{W}(\text{concat}([E^{(1)}, E^{(2)}, E^{int}]))$ | 0.474 | 0.694 |
| $\mathbf{W}(\text{BatchNorm}(\text{concat}([E^{(1)}, E^{(2)}, E^{int}])))$ | 0.378 | 0.670 |

**Table A2.**
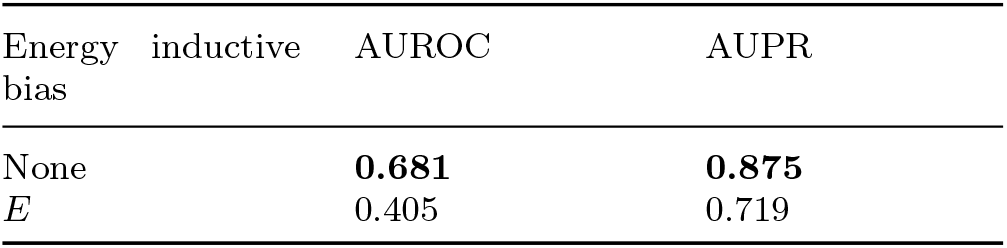
Energy inductive bias doesn’t help with the ProtCID task.

**Table A3.**
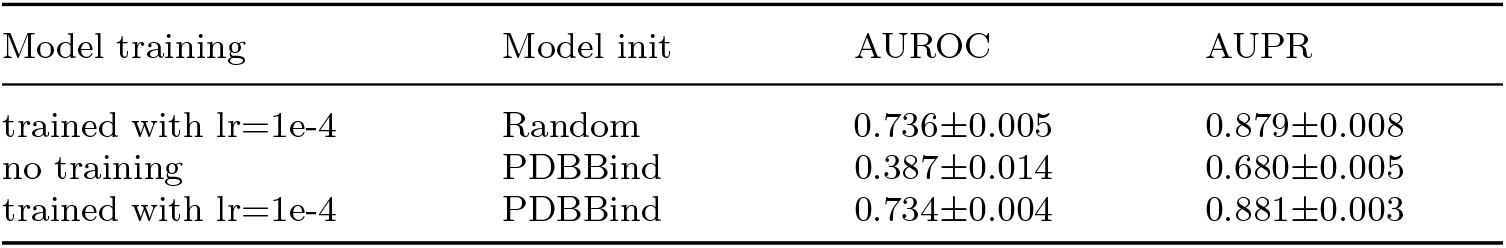
Transferring model weights learned from PDBBind to ProtCID. Experiments were performed using GIN as the lower-level GNN within the single-stage EGGNet architecture. Average and standard deviations are shown in the table from 3 independent runs with different random seeds. No significant differences (t-test p-value *>*0.6) were observed between transfer learning and training from scratch.

**Fig. A1.**
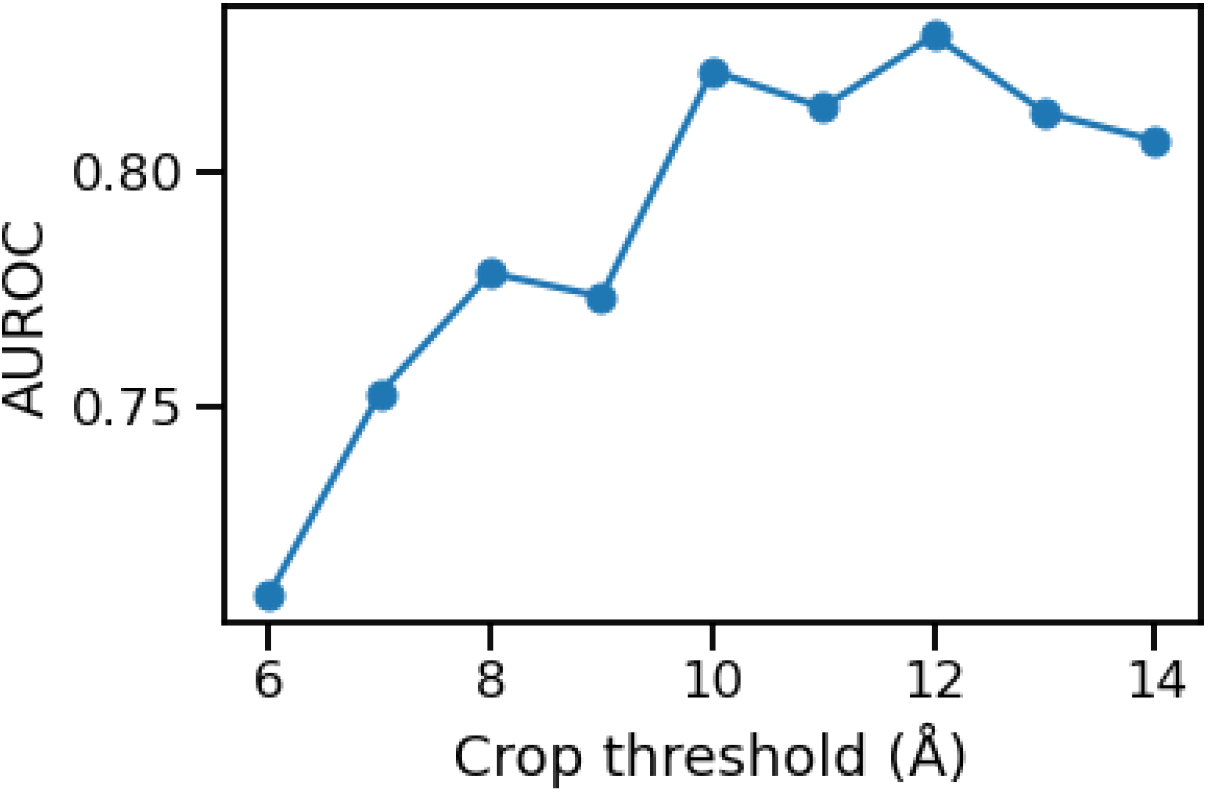
Performance of EGGNet on the biological versus crystal interfaces classification task from the DC dataset. AUROC are shown across a range of cropping thresholds used to isolate the protein-protein interfaces when constructing the GoGs. Experiments were performed using GIN as the lower-level GNN within the single-stage EGGNet architecture without joint training. We found 12Å to be the optimal threshold for isolating the interface.

